# Repression of LKB1 by *miR-17∼92* sensitizes *MYC*-dependent lymphoma to biguanide treatment

**DOI:** 10.1101/2019.12.20.883025

**Authors:** Said Izreig, Alexandra Gariepy, Ariel O. Donayo, Gaëlle Bridon, Daina Avizonis, Ryan D. Sheldon, Rob Laister, Mark D. Minden, Nathalie A. Johnson, Kelsey S. Williams, Marc S. Rudoltz, Thomas F. Duchaine, Sanghee Yoo, Michael N. Pollak, Russell G. Jones

**Author notes:** Corresponding Author: Russell G. Jones, Metabolic and Nutritional Programming, Center Cancer and Cell Biology, Van Andel Research Institute.

## Abstract

Cancer cells display metabolic plasticity to survive metabolic and energetic stresses in the tumor microenvironment, prompting the need for tools to target tumor metabolism. Cellular adaptation to energetic stress is coordinated in part by signaling through the Liver Kinase B1 (LKB1)-AMP-activated protein kinase (AMPK) pathway. Reducing LKB1-AMPK signaling exposes metabolic vulnerabilities in tumor cells with potential for therapeutic targeting. Here we describe that miRNA-mediated silencing of LKB1 (mediated by the oncogenic miRNA cluster *miR-17∼92*) confers sensitivity of lymphoma cells to mitochondrial inhibition by biguanides. Using both classic (phenformin) and novel (IM156) biguanides, we demonstrate that *Myc*^+^ lymphoma cells with elevated *miR-17∼92* expression display increased sensitivity to biguanide treatment both in cell viability assays *in vitro* and tumor growth assays *in vivo*. This increased biguanide sensitivity is driven by *miR-17*-dependent silencing of LKB1, which results in reduced AMPK activation in response to bioenergetic stress. Mechanistically, biguanide treatment inhibits TCA cycle metabolism and mitochondrial respiration in *miR-17∼92*-expressing tumor cells, targeting their metabolic vulnerability. Finally, we demonstrate a direct correlation between *miR-17∼92* expression and biguanide sensitivity in human cancer cells. Our results identify *miR-17∼92* expression as a potential biomarker for biguanide sensitivity in hematological malignancies and solid tumors.

**One Sentence Summary:** miR-17∼92 expression in *Myc*^*+*^ tumors sensitizes cancer cells to biguanide treatment by disrupting bioenergetic stability in lymphoma cells.

## Introduction

A major driver of metabolic reprogramming in cancer is the c-Myc proto-oncogene (Myc), a transcription factor that broadly regulates the expression of genes involved in anabolic metabolism and cellular bioenergetics *(1)*. Oncogene activation or tumor suppressor inactivation promotes changes in cellular metabolism that facilitate transformation and tumor cell proliferation *(2)*. However, the increased metabolic activity stimulated by transformation imposes additional stresses on tumor cells such as nutrient depletion and redox imbalance, which must be countered in order for tumors to survive and grow. The Liver Kinase B1 (LKB1)/AMP-activated protein kinase (AMPK) pathway contributes to tumor cell survival by promoting cellular sensing of and adaptation to such bioenergetic stress *(3)*. Inactivation of LKB1 in cancer, conversely, promotes anabolic metabolic reprogramming at the expense of metabolic flexibility *(4–7)*. In cells lacking LKB1, including non-small cell lung cancer (NSCLC), AMPK activation is reduced and cells are more sensitive to metabolic stresses induced by nutrient limitation or inhibitors of oxidative phosphorylation (OXPHOS) *(8, 9)*. This raises the possibility of using OXPHOS inhibitors to target cancers with reduced bioenergetic capacity and/or defects in energy-sensing control systems. Indeed, potent OXPHOS inhibitors (for example IACS-010759) *(10)*, putative inhibitors of mitochondrial protein synthesis *(11, 12)*, and enhancers of degradation of respiratory chain proteins *(13, 14)* are all under study as antineoplastic strategies.

Biguanides represent another class of drugs proposed to target cellular bioenergetics in tumor cells. The most well-known biguanide is metformin—a drug commonly used to treat type II diabetes. Metformin reduces the activity of mitochondrial complex I, thereby limiting mitochondrial ATP production and imposing bioenergetic stress on cells *(15, 16)*. This leads to reduced hepatic gluconeogenesis and reduced hyperglycemia in type II diabetes. However, if adequate drug levels can be achieved in other cell types, lineage-specific consequences of energetic stress are observed, including cytostatic or cytotoxic effects on cancer cells *(17)*. Older retrospective studies *(18)* reported that diabetics treated with metformin had reduced cancer risk and better cancer prognosis relative to diabetic patients not taking metformin *(18)*, generating interest in repurposing biguanides as anti-cancer agents *(17)*. However, further pharmaco-epidemiologic studies *(19, 20)* failed to confirm these findings, and early randomized clinical trials of metformin in advanced cancer have shown no survival benefit in pancreatic cancer *(21)*, and only minor benefit in lung cancer *(22)*. Pharmacokinetic factors, such as the requirement for active transport of metformin by the organic cation 1 (OCT1) transporters, may account in part to the discrepancies between preclinical models and clinical trials with respect to anti-cancer effects of metformin. Nevertheless, many *in vivo* cancer models demonstrate significant *in vivo* antineoplastic activity of biguanides *(6, 23–27)*, raising the possibility that biguanides with better bioavailability and toxicity profiles may have clinical utility.

Important in the clinical development of OXPHOS inhibitors as antineoplastic drugs is selection of subsets of cancers that are particularly sensitive to metabolic stress. Preclinical work by Shackelford *et al.* demonstrated that biguanides, specifically phenformin, could be effective as single agents for LKB1-deficient KRAS-mutant NSCLC *(8)*, in keeping with LKB1’s role in adaptation to energetic stress. While mutation of LKB1 is found in ∼20–30% of NSCLC, we hypothesized that biguanide-sensitive cancers can be extended to those with increased expression of MYC, which we have previously reported promotes translational suppression of LKB1 via the miRNA *miR-17∼92 (28)*. In this study, using loss- and gain-of-function models of *miR-17∼92*, we demonstrate that elevated *miR-17∼92* expression, specifically the seed family *miR17/20*, confers increased sensitivity of mouse and human lymphoma cells to apoptosis induced by two biguanides—phenformin and the novel biguanide IM156. As single agents, both phenformin and IM156, but not metformin, extended the survival of mice bearing *miR-17∼92*-expressing lymphomas. Collectively, these results suggest that *miR-17∼92* could potentially function as a biomarker for biguanide sensitivity in lymphomas.

## Results

### IM156 is a novel biguanide that inhibits mitochondrial respiration

The limited bioavailability of metformin and its dependence on OCT1 for cellular uptake potentially limits its applicability in the treatment of cancer *(29)*. We investigated the biological properties of phenformin and the novel biguanide IM156, which are more hydrophobic and therefore potentially more bioavailable to cells than metformin (**Figure 1A**). To test the impact of these biguanides on tumor cell respiration, we acutely treated *Myc*-dependent mouse lymphoma cells (*Eµ-Myc* cells) with either metformin, phenformin, or IM156 and assessed changes in oxygen consumption rate (OCR) using the Seahorse XF96 extracellular flux analyzer. Across a range of concentrations, phenformin and IM156 decreased OCR (**Figures 1B**), with IM156 exhibiting greater potency than phenformin and metformin at equal concentrations. IM156 was more effective than phenformin at reducing cellular ATP production at equal concentrations, correlating with the effect of IM156 on oxidative phosphorylation (**Figure 1C**). These data are consistent with IM156 functioning as a more potent inhibitor of mitochondrial respiration than phenformin.

**Fig. 1.**
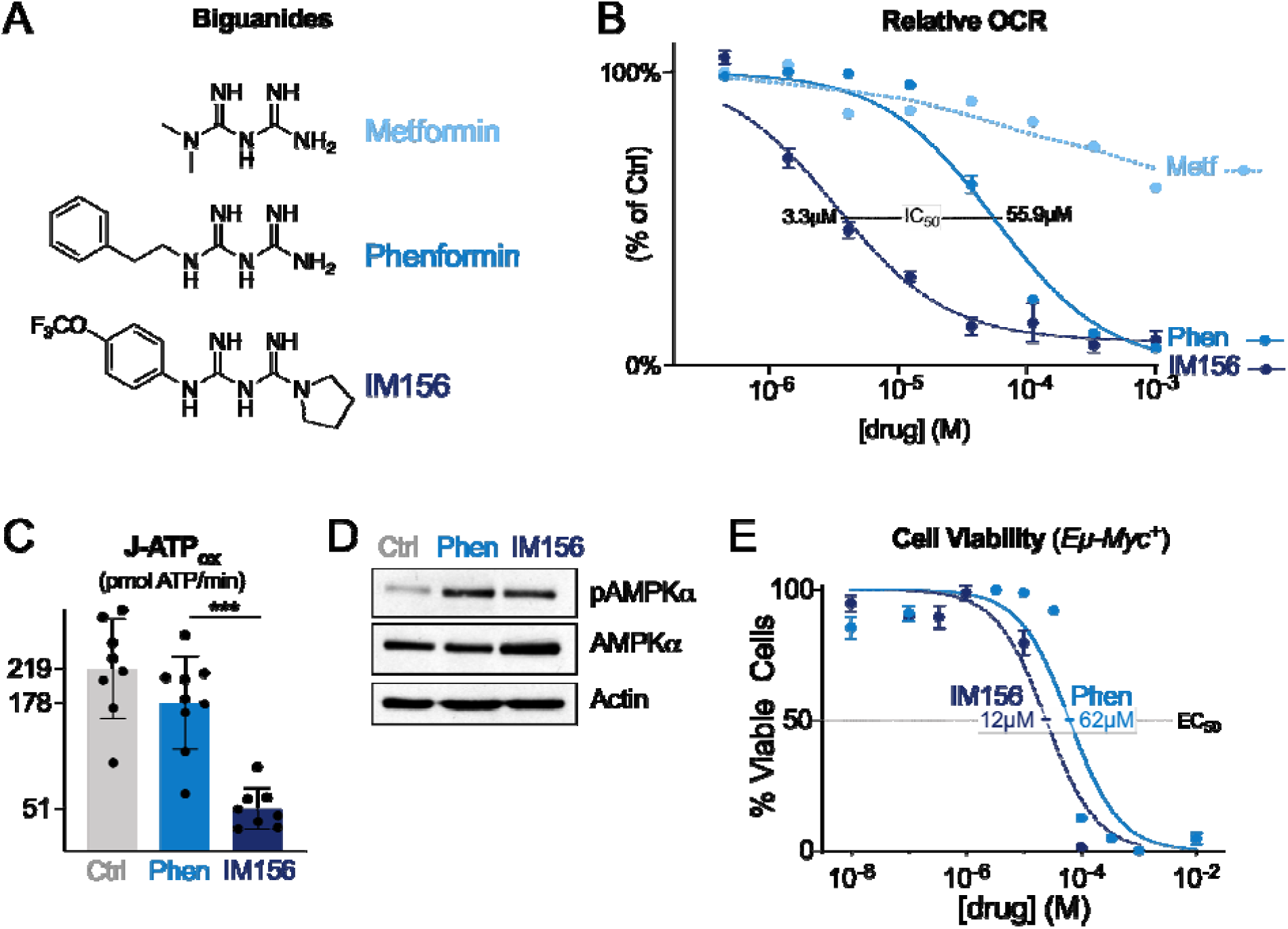
IM156 is a novel biguanide that inhibits mitochondrial respiration. **(A)** Chemical structure of the biguanides metformin, phenformin, and IM156. **(B)** Dose-dependent reduction of OCR of Eµ-*Myc*^+^ lymphoma cells by metformin, phenformin, and IM156. Data were derived from the maximal point of OCR reduction after 30 minutes of treatment with the indicated biguanide and expressed relative to untreated cells at the same timepoint. Data represent mean ± SEM for biological replicates (n = 4–6 per drug per concentration). **(C)** Mitochondrial ATP production rate of Eµ-*Myc*^+^ lymphoma cells after incubation with 100 µM of either vehicle, phenformin (Phen), or IM156. Data represent mean ± SD for biological replicates (n = 8–9 per group). **(D)** Immunoblot for phosphorylated (pAMPK, T172) and total AMPKα following treatment with vehicle or equivalent doses of phenformin (100 µM) or IM156 (10 µM) for 2 hours. Actin is shown as a loading control. **(E)** Viability of Eµ-*Myc*^+^ lymphoma cells following 48 h treatment with phenformin or IM156, comparing ten different doses. EC_50_ for each treatment is indicated. Data represent mean ± SEM for biological replicates (n = 3 per drug per concentration). Statistics for all figures are as follows: *, *p<0.05*; **, *p<0.01*; ***, *p<0.001*.

We next determined if phenformin and IM156 similarly activate the energy sensor AMPK, as depletion of cellular ATP by biguanide treatment is a known trigger for AMPK activation *(30)*. Given the difference in potency of phenformin and IM156, we used doses (100 µM and 10 µM, respectively) that produced an equivalent ∼50% decrease in OCR from baseline after two hours treatment (**Figure S1A–B**). Two-hour treatment with these concentrations of phenformin or IM156 yielded similar levels of AMPK phosphorylation (**Figure 1D**). Lastly, we treated *Eµ-Myc*^+^ lymphoma cells with a range of concentrations of both phenformin and IM156. Based on cell viability measurements, IM156 exhibited higher potency and induced lymphoma cell death at lower concentrations than phenformin (EC_50_ of 12 µM for IM156 compared to 62 µM for phenformin, **Figure 1E**).

### miR-17∼92 *sensitizes lymphoma cells to apoptosis by biguanides*

We previously demonstrated that the oncogenic miRNA cluster *miR-17∼92* is required for *Myc*-dependent metabolic reprogramming in lymphoma and does so in part through translational suppression of LKB1 *(28)*. We next examined whether expression of *miR-17∼92* alters the sensitivity of lymphoma cells to biguanide treatment. We used Eµ-*Myc* B cell lymphoma cells harboring floxed *miR-17∼92* alleles, which allowed us to study the effect of conditional deletion of *miR-17∼92* in the presence of constitutive *Myc* expression *(31)*. Eµ-*Myc* lymphoma cells deleted for *miR-17∼92* (Δ/Δ) were more resistant to phenformin treatment than their isogenic counterparts expressing *miR-17∼92* (*fl/fl*) (**Figure 2A**). Phenformin treatment actively induced apoptosis in *miR-17∼92*-expressing Eµ-*Myc* lymphoma cells as shown by the presence of active (cleaved) caspase-3 (**Figure 2B**). Levels of caspase-3 cleavage were markedly reduced in Eµ-*Myc* lymphoma cells lacking *miR-17∼92* (**Figure 2B**).

**Fig. 2.**
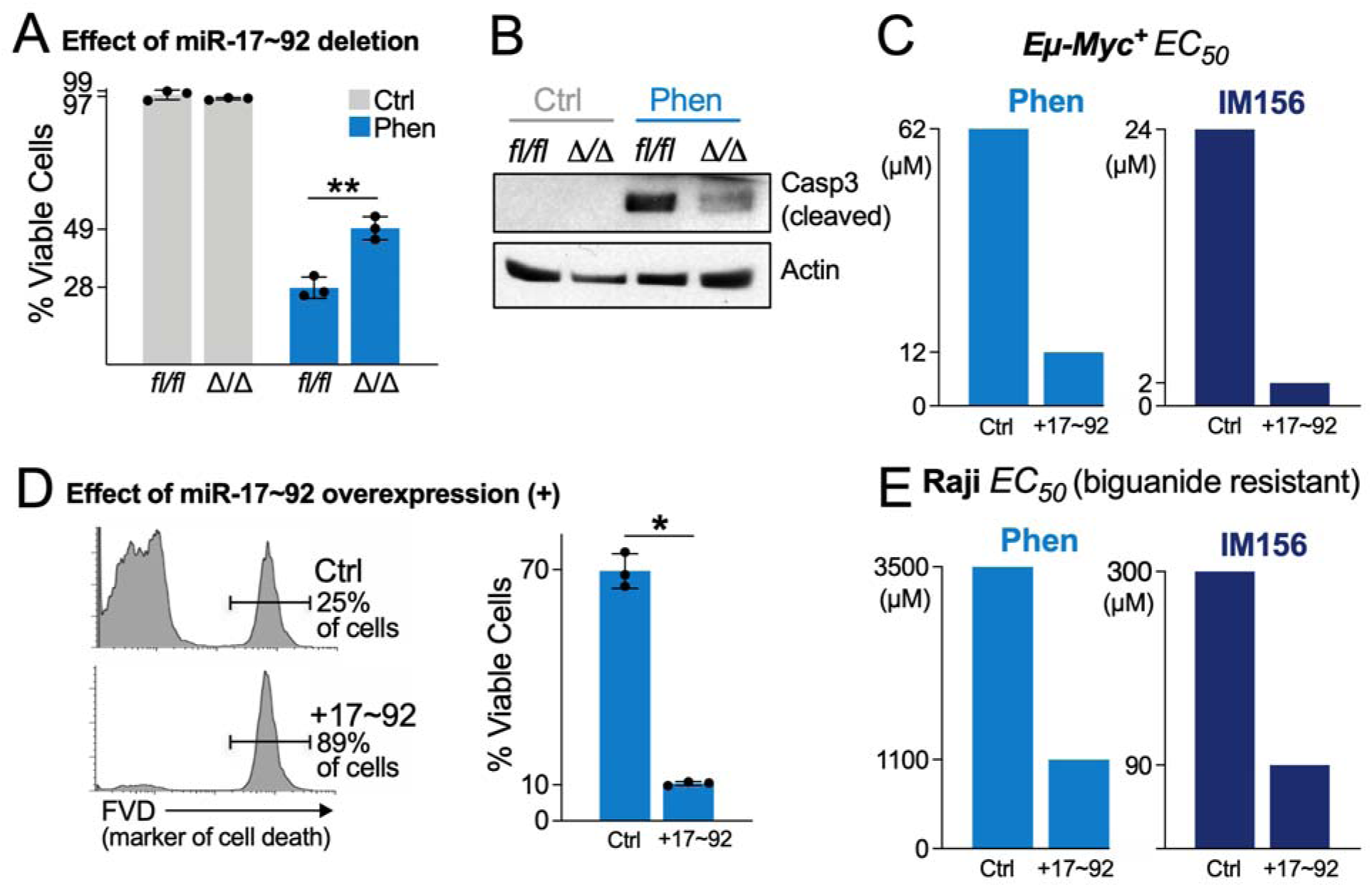
*miR-17∼92* sensitizes lymphoma cells to apoptosis by biguanides. **(A)** Viability of Ctrl (fl/fl) and miR-17∼92-deficient (Δ/Δ) Eµ-*Myc*^+^ lymphoma cells untreated (Ctrl, grey) or treated with 100 μM phenformin (Phen, blue) for 48 h. Data represent mean ± SD for biological replicates (n = 3 per group). **(B)** Immunoblot for active (cleaved) caspase-3 in cells treated as in (A). **(C)** Dose of phenformin (left) or IM156 (right) required to achieve 50% decrease in cell viability (EC_50_) in biguanide sensitive Eµ-*Myc*^+^ lymphoma cells expressing control (Ctrl) or miR-17∼92 (+17∼92) expression vectors. Cell viability was measured 48 h post biguanide treatment. **(D)** Viability of control (Ctrl) or miR-17∼92-expressing (+17∼92) Raji cells after 48 h treatment with 100 μM phenformin. Histogram shows representative staining for cell death using fluorescent viability dye (FVD), which is quantified on the right. Data represent mean ± SD for biological replicates (n = 3 per group). **(E)** Dose of phenformin (left) or IM156 (right) required to achieve 50% decrease in cell viability (EC_50_) in biguanide-resistant Raji cells expressing control (Ctrl) or miR-17∼92 (+17∼92) expression vectors following 48 h treatment with biguanide. Statistics for all figures are as follows: *, *p<0.05*; **, *p<0.01*; ***, *p<0.001*.

Since *miR-17∼92* is recurrently amplified in lymphoma *(32, 33)*, we next tested whether increased copy number of *miR-17∼92* was sufficient to increase the sensitivity of lymphoma cells to biguanides. To test this, we generated Eµ-*Myc* lymphoma cells and Raji lymphoma cells—a human Burkitt’s lymphoma cell line known to display low Myc levels *(28)*—with ectopic expression of the entire *miR-17∼92* polycistron (hereafter denoted as +*17∼92*). *Eµ-Myc* lymphoma cells overexpressing *miR-17∼92* were significantly more sensitive than control cells when treated with either phenformin or IM156 (**Figure 2C** and **S1A**). *miR-17∼92* overexpression led to a 10-fold shift in the EC_50_ of *Eµ-Myc* cells to IM156 treatment (2 μM versus 24 μM). Similar results were observed in Raji cells engineered to express higher levels miR-17∼92 (**Figure 2D–E** and **S1B**). Although Raji cells were less sensitive to biguanide treatment, with EC_50_ ranging from 90–3500 µM as compared to *Eµ-Myc*^+^ EC_50_ ranging from 2–62 µM (**Figure S1A–B**), overexpression of *miR-17∼92* still conferred a ∼3-fold increase in sensitivity to biguanides compared to control cells. These data indicate that elevated *miR-17∼92* enhances sensitivity of lymphoma cells to biguanide treatments, and that *Myc* and *miR-17∼92* synergize to increase sensitivity to biguanide treatment.

We next asked whether the enhanced sensitivity to biguanides observed in vitro extended to in vivo models. Nude mice harboring either control or *miR-17∼92*-overexpressing *Eµ-Myc* cells were administered phenformin or IM156 in their drinking water *ad libitum* and survival was tracked compared to tumor-bearing mice administered regular drinking water. While biguanide treatment of mice bearing control lymphoma cells produced no discernable survival benefit (**Figure 3A**), both phenformin and IM156 significantly prolonged the lifespan of mice bearing aggressive *miR-17∼92*-overexpressing tumors (**Figure 3B**). Our observations indicate that phenformin and IM156 can act as single agents to extend survival of mice bearing tumors with elevated *miR-17∼92* expression. Together these data indicate that elevated *miR-17∼92* expression is a predictor of sensitivity to biguanides.

**Fig. 3.**
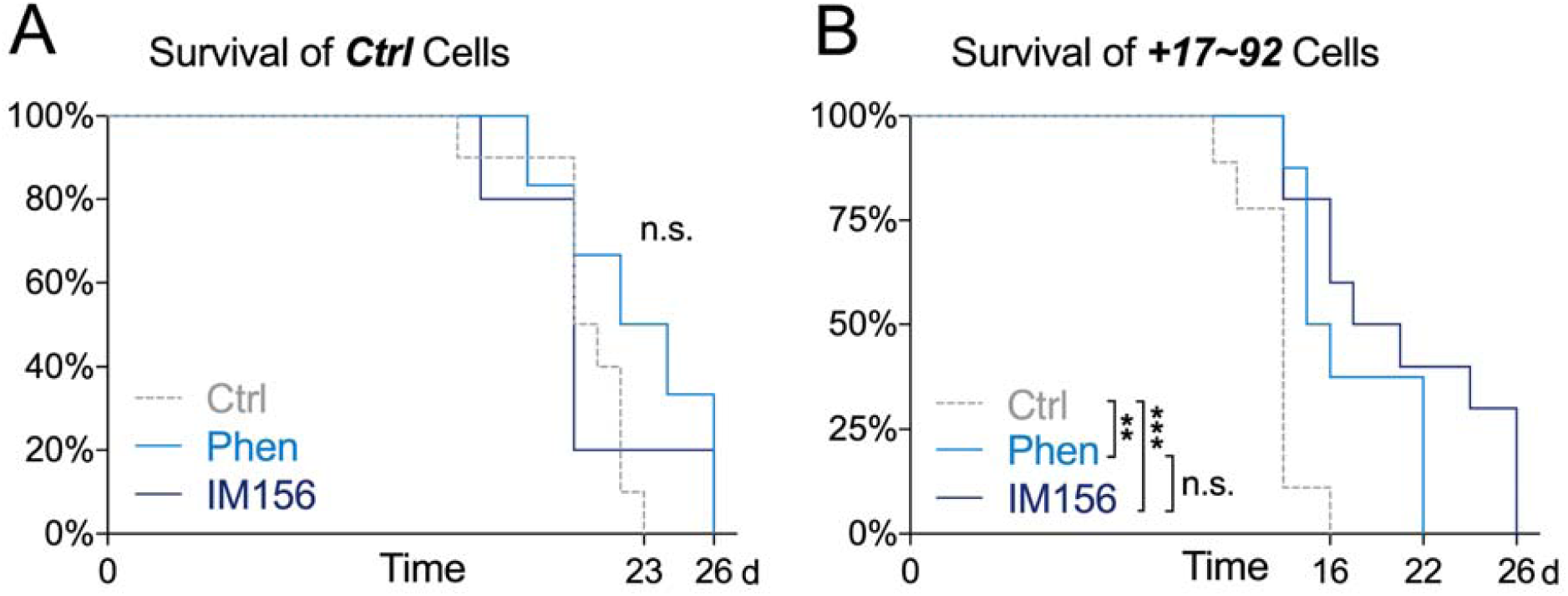
Biguanide treatment selectively impairs the growth of *miR-17∼92*-expressing lymphoma cells *in vivo*. **(A–B)** Kaplan-Meier curves for viability of nude mice injected with 1×10^6^ *Myc*^+^/Ctrl (A) or *Myc*^+^/+17∼92 (B) lymphoma cells. Mice were provided with untreated water (Ctrl, n = 10), 0.9 mg/mL phenformin (Phen, n = 8), or 0.8 mg/mL IM156 (IM156, n = 10) *ad lib* following intravenous tumor cell injection. Statistics for all figures are as follows: *, *p<0.05*; **, *p<0.01*; ***, *p<0.001*.

### miR-17∼92 *confers sensitivity to biguanides through suppression of LKB1*

We next explored the mechanism by which elevated miR-17∼92 expression confers biguanide sensitivity in lymphoma cells. Non-small cell lung cancer (NSCLC) cells that have lost LKB1 display increased sensitivity to phenformin *(8)*. We previously identified LKB1 as a target of gene silencing by miR-17∼92 via miR-17- and 20-dependent suppression of LKB1 protein expression *(28)*. Deletion of *miR-17∼92* in *Eµ-Myc* cells (Δ/Δ cells) increased LKB1 protein levels (**Figure 4A**), while expression of *miR-17∼92* reduced LKB1 expression (**Figure 4B**), resulting in reduced AMPK phosphorylation (**Figure 4B**) and enhanced mTORC1 signaling (**Figure 4C**). Deletion of *miR-17∼92* in *Eµ-Myc* cells (Δ/Δ cells) was sufficient to confer a survival advantage in response to IM156 treatment (**Figure 4D**). Reducing LKB1 expression in Δ/Δ cells using shRNAs targeting LKB1*(28)* restored biguanide sensitivity in Δ/Δ cells to match that observed in lymphoma cells expressing *miR-17∼92* (**Figures 4D**). Similarly, lymphoma cells expressing *miR-17∼92* but lacking expression of miR-17 were resistant to IM156 treatment (**Figure 4E**).

**Fig. 4.**
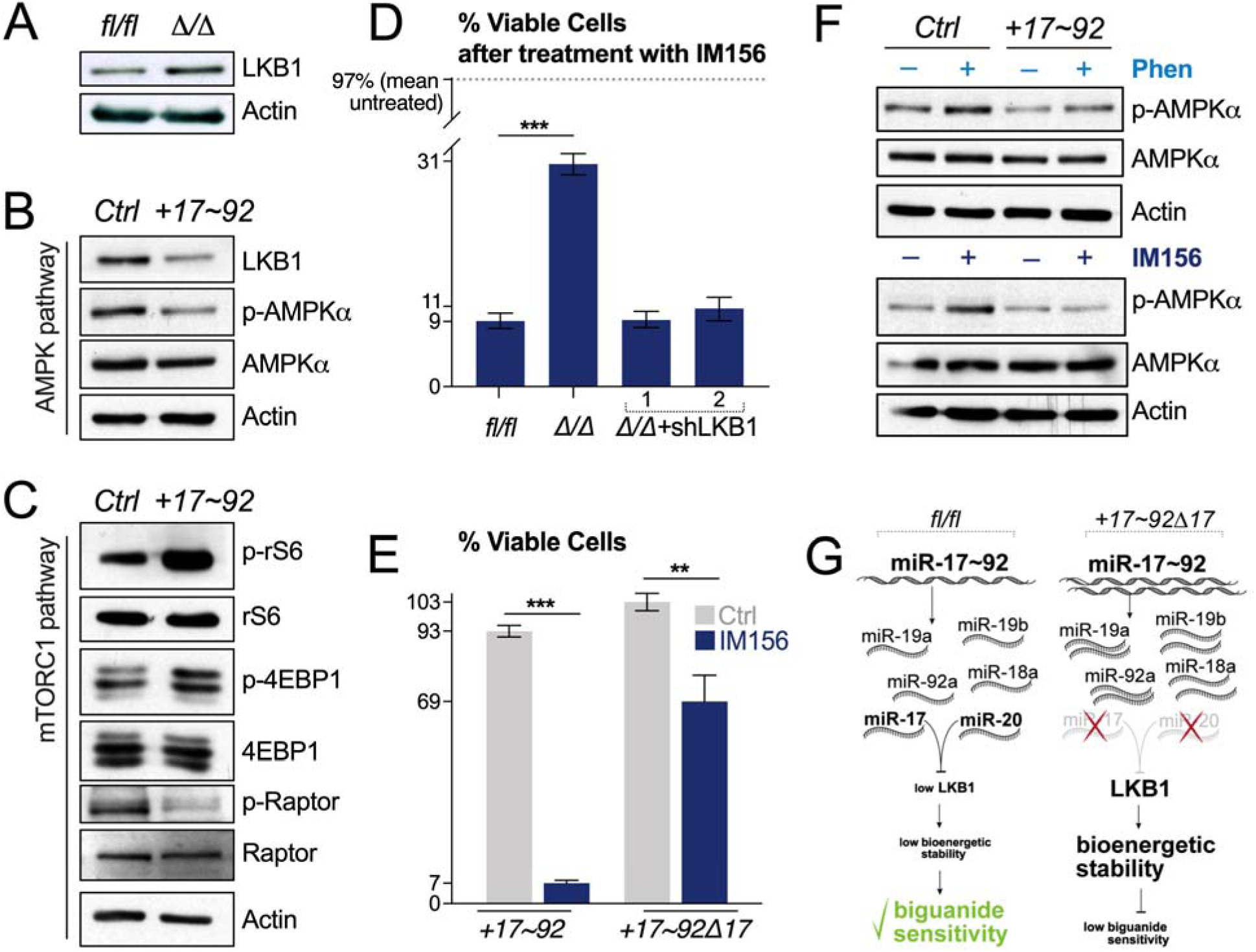
*miR-17∼92* confers sensitivity to biguanides through suppression of LKB1. **(A)** Immunoblot of LKB1 and actin protein levels in Ctrl (fl/fl) and miR-17∼92-deficient (Δ/Δ) Eµ-*Myc*^+^ lymphoma cells. **(B–C)** Immunoblot of AMPK pathway (B) and mTORC1 pathway (C) activation in control (Ctrl) or miR-17∼92-expressing (+17∼92) Eµ-*Myc*^+^ lymphoma cells. AMPK activation was determined by measuring total and phosphorylated (p-AMPK, T172) AMPKα. mTORC1 activity was assessed by measuring levels of total and phosphorylated ribosomal S6 protein (rS6), 4E-BP, and Raptor. **(D)** Viability of the following Eµ-*Myc*^+^ lymphoma cells: Ctrl (fl/fl), miR-17∼92-deficient (Δ/Δ), miR-17∼92-deficient expressing shRNAs targeting LKB1 (Δ/Δ+shLKB1/2). Cells were treated with 10 μM IM156 for 48 h and viability assessed by flow cytometry. Data represent mean ± SD for biological replicates (n = 3 per group). **(E)** Viability of Eµ-*Myc*^+^ lymphoma cells expressing miR-17∼92 (+17∼92) or miR-17∼92 lacking the miR-17 and 20 seed families (+17∼92Δ17). Cells were treated with vehicle (Ctrl) or 10 μM IM156 for 48 h and viability assessed by flow cytometry. Data represent mean ± SD for biological replicates (n = 3 per group). **(F)** Immunoblot of total and phosphorylated (p-AMPK, T172) AMPKα levels in control (Ctrl) and miR-17∼92-expressing (+17∼92) Eµ-*Myc*^+^ lymphoma cells following 2 h treatment with vehicle (–), 100 µM phenformin (+ Phen), or 10 μM IM156 (+ IM156). **(G)** Schematic of transcriptional differences in miRNA between control (fl/fl) and miR-17∼92 expressing Eµ-*Myc*^+^ lymphoma cells. miR-17 and -20 are responsible for repression of LKB1, which increases biguanide sensitivity. Statistics for all figures are as follows: *, *p<0.05*; **, *p<0.01*; ***, *p<0.001*.

Given that an increased copy number of *miR-17∼92* leads to LKB1 repression in lymphoma cells (**Figure 4B**), we tested whether AMPK activation was differentially engaged downstream of either phenformin or IM156 treatment. Lymphoma cells overexpressing *miR-17∼92* displayed reduced AMPK phosphorylation following phenformin or IM156 treatment compared to control cells (**Figure 4F**), suggesting uncoupling of the LKB1-AMPK axis to metabolic stress (**Figure 4G**).

### *Biguanide treatment affects central carbon metabolism in* miR-17∼92-*expressing Myc*^+^ *lymphoma cells*

We next characterized the effect of IM156 treatment on lymphoma cell bioenergetics using a Seahorse XF96 extracellular flux analyzer. Lymphoma cells overexpressing *miR-17∼92* exhibit higher basal metabolic rate—both in terms of glycolytic rate (ECAR) and respiration (OCR)—than control cells (**Figure 5A**). This corresponded to higher overall rates of ATP production by *miR-17∼92*-expressing lymphoma cells compared to controls, which was due largely to increased glycolytic ATP production (**Figure 5B**). *miR-17∼92*-expressing lymphoma cells also displayed a larger bioenergetic scope than control cells (**Figure 5C**). The bioenergetic profile of both cell types was significantly altered by IM156 treatment, characterized by a drop in mitochondrial ATP production (*J*_*ATP*_ *ox*) and slight increase in glycolytic ATP production (*J*_*ATP*_ *gly*) (**Figure 5C**). IM156 reduced OXPHOS-derived ATP production in both cell types by greater than 50% (**Figure 5C**).

**Fig. 5.**
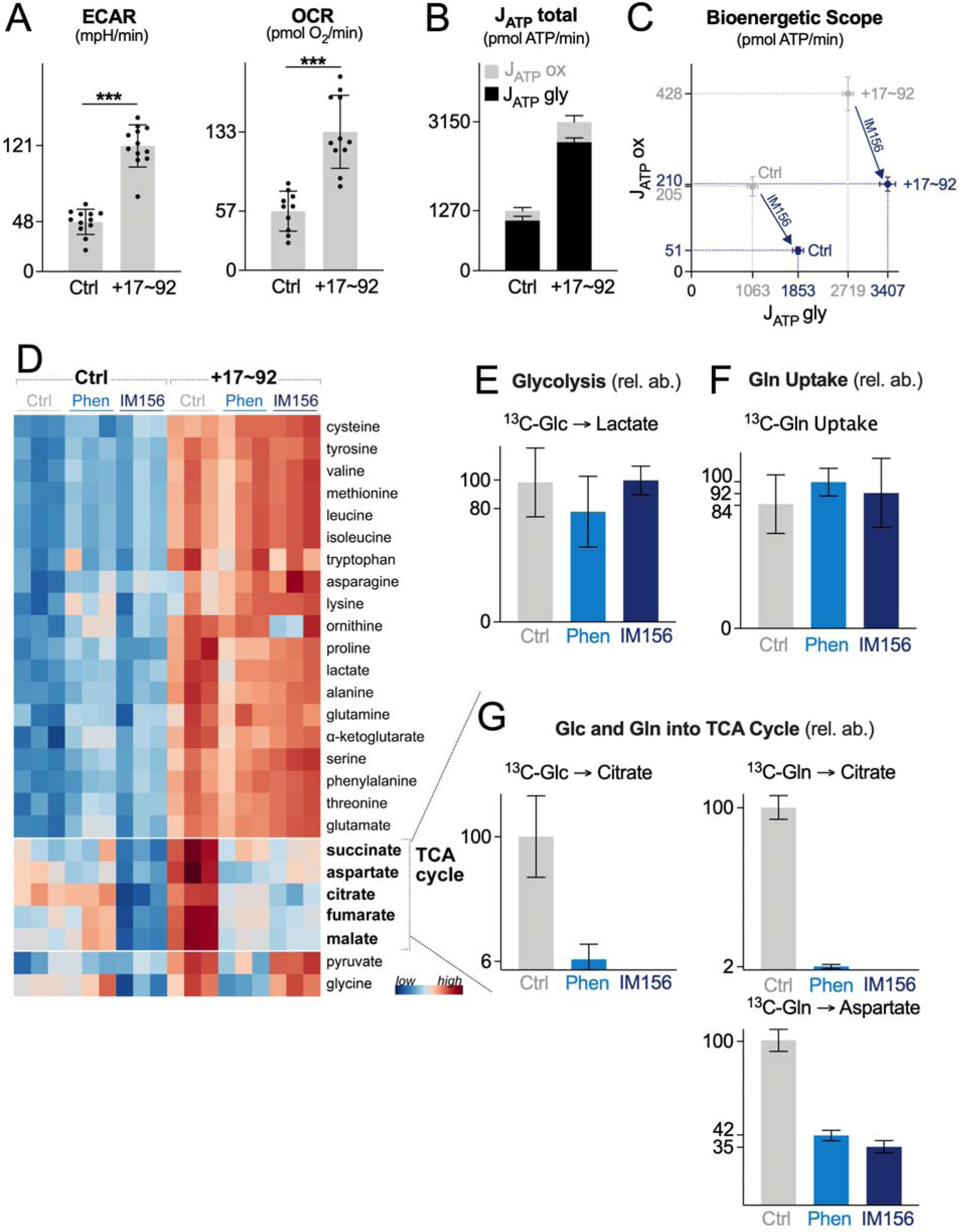
Biguanide treatment affects central carbon metabolism in *miR-17∼92*-expressing lymphoma cells. **(A)** ECAR and OCR of control (Ctrl) or miR-17∼92-expressing (+17∼92) Eµ-*Myc*^+^ lymphoma cells. Data represent mean ± SD for biological replicates (n = 12 per group). **(B)** ATP production rates (J_ATP_) for Eµ-*Myc*^+^ lymphoma cells expressing control (Ctrl) or miR-17∼92 (+17∼92) expression vectors. Seahorse experiments were performed under standard cell culture media conditions (Glc, 25 mM; Gln, 2 mM). J_ATP_total is the sum of the glycolytic (J_ATP_gly) and OXPHOS (J_ATP_ox) ATP production rates. Data represent mean ± SD for biological replicates (n = 12 per group). **(C)** Bioenergetic capacity plot of Eµ-*Myc*^+^ lymphoma cells expressing control (Ctrl) or miR-17∼92 overexpression (+17∼92) vectors following 2 h treatment with vehicle (gray) or IM156 (purple). Rectangles define the maximum bioenergetic space of each cell type. Data represent mean ± SEM for biological replicates (Ctrl, n = 8; Ctrl+IM156, n = 8; +17∼92, n = 11; +17∼92+IM156, n = 12). **(D)** Heatmap showing relative metabolite abundances for control (Ctrl) or miR-17∼92-expressing (+17∼92) Eµ-*Myc*^+^ lymphoma cells following 2 h treatment with vehicle (Ctrl), phenformin (Phen), or IM156. (n = 3 per group) **(E)** Fractional enrichment of U-[^13^C]-glucose-derived lactate in miR-17∼92-expressing Eµ-*Myc*^+^ lymphoma cells following 2 h treatment with vehicle (Ctrl), phenformin (Phen), or IM156. Data represent mean ± SD for biological replicates (n = 3 per group). **(F)** Fractional enrichment of ^13^C-labelled glutamine in miR-17∼92-expressing Eµ-*Myc*^+^ lymphoma cells following 2 h treatment with vehicle (Ctrl), phenformin (Phen), or IM156. Data represent mean ± SD for biological replicates (n = 3 per group). **(G)** Fractional enrichment of U-[^13^C]-glucose-derived citrate and ^13^C-glutamine-derived citrate and aspartate in miR-17∼92-expressing Eµ-*Myc*^+^ lymphoma cells, following 2 h treatment with vehicle (Ctrl), phenformin (Phen), or IM156. Data represent mean ± SD for biological replicates (n = 3 per group). Statistics for all figures are as follows: *, *p<0.05*; **, *p<0.01*; ***, *p<0.001*.

We next examined the impact of biguanide treatment on metabolite dynamics in lymphoma cells. Consistent with their increased bioenergetic profile (**Figure 5A–C**), *miR-17∼92*-expressing lymphoma cells displayed increased abundance of metabolites involved in glycolysis (e.g., pyruvate, lactate), TCA cycle (e.g., citrate, succinate, fumarate, malate), and amino acid metabolism (**Figure 5D**). Biguanide treatment had minimal effect on amino acid levels in either cell type (**Figure 5D**). However, levels of TCA cycle intermediates were markedly reduced in *miR-17∼92*-expressing cells upon treatment with phenformin or IM156 (**Figure 5D**), suggesting that biguanide treatment limits metabolite flux through the TCA cycle. This was not due to an inhibition of glucose or glutamine uptake, as ^13^C-tracing experiments revealed no change in glucose trafficking to lactate (**Figure 5E**) or glutamine uptake (**Figure 5F**) in the presence of phenformin or IM156. Biguanide treatment blunted ^13^C-glucose and ^13^C-glutamine entry into the TCA cycle, as evidenced by reduced conversion of glucose and glutamine to citrate (**Figure 5G**). ^13^C-glutamine-derived production of aspartate, a key intermediate for protein and nucleotide biosynthesis and tumor cell growth *(34, 35)*, was reduced by 60–65% following 2 h administration of phenformin or IM156. These data suggest that although *miR-17∼92* enhances central carbon metabolism and bioenergetic potential in lymphoma cells, these pathways are uniquely sensitive to biguanide treatment.

### miR-17/20 *expression correlates with biguanide sensitivity in human lymphoma cells*

Having defined the downstream effects of biguanides on the metabolism and viability of *miR-17∼92*-expressing *Myc*^+^ lymphoma cells, we expanded our analysis to test biguanide sensitivity in a panel of human lymphoma cell lines. Ten verified *Myc*^+^ lymphoma cell lines were used for this analysis: SU-DHL-4, OCI-Ly7, Jeko-1, Rec-1, Karpas 1718, OCI-Ly-1, OCI-Ly-2, OCI-Ly-3, OCI-Ly-8, and OCI-Ly-18. For each cell line we measured levels of both the primary (*pri-miR*) and mature (*miR*) forms of *miR-17* and *miR-20* and determined the EC_50_ for cell viability to phenformin or IM156. From these data, we observed a negative correlation between the relative level of *miR-17/20* expressed in cells and the dose of biguanide required to achieve EC_50_ (**Figure 6A–B** and **S2**), with cells displaying the highest *miR-17/20* expression showing the greatest sensitivity to biguanide treatment. Levels of *pri-miR-17* were a better predictor of biguanide sensitivity than mature *miR-17* (**Figure 6A–B**). IM156 was more potent than phenformin in inducing cell death in human lymphoma cells, similar to that observed in *Eµ-Myc*^+^ lymphoma cells. Together these data demonstrate *miR-17* expression levels to be a biomarker for biguanide sensitivity.

**Fig. 6.**
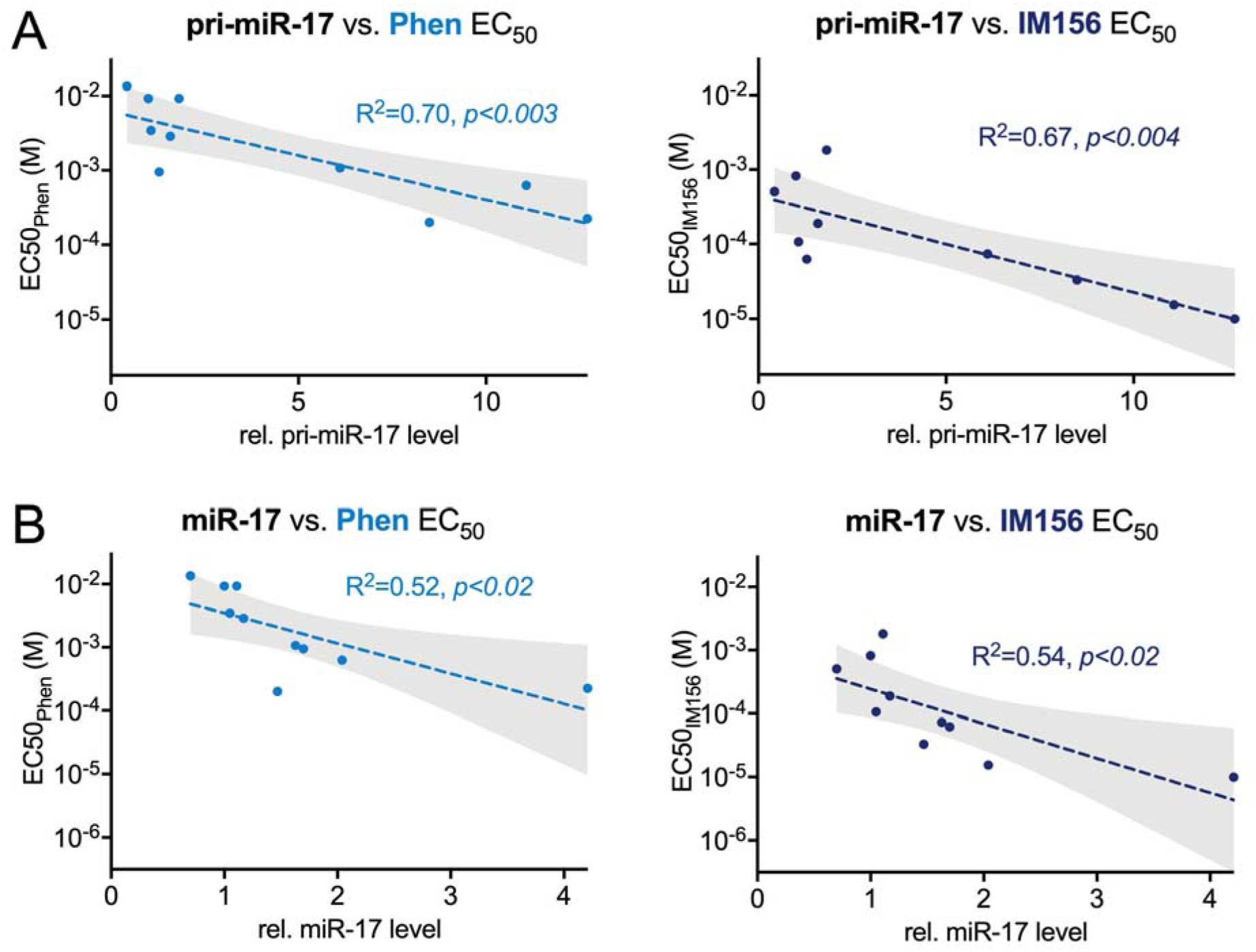
*miR-17/20* expression correlates with biguanide sensitivity in human lymphoma cells. **(A–B)** EC_50_ for phenformin (Phen, left) and IM156 (right) correlated to the relative levels of pri-miR-17 (A) and mature miR-17 (B) transcripts for ten human lymphoma cell lines. Linear regression (dotted line) is shown for both drugs with shaded 95% confidence interval. Statistics for all figures are as follows: *, *p<0.05*; **, *p<0.01*; ***, *p<0.001*.

## Discussion

Myc is abnormally expressed in many human cancers; however, development of small molecule Myc inhibitors has been a persistent challenge *(36)*, prompting consideration of alternative approaches to *Myc* targeting. Here we report that a downstream target of *Myc*-driven metabolism—*miR-17∼92*—while increasing bioenergetic capacity of tumor cells, also increases sensitivity to apoptosis induced by biguanides. Using both classic (phenformin) and novel (IM156) biguanides, we demonstrated that human and mouse lymphoma cells with elevated *miR-17∼92* expression are highly sensitized to biguanide treatment. As single agents, both phenformin and IM156 were effective at extending survival of mice bearing *miR-17∼92*-expressing lymphomas. In addition, we observed a statistical correlation between *miR-17* levels in human lymphoma cells and sensitivity to biguanide treatment. Together our data provide rationale for the clinical investigation of miR-17∼92 expression level as a potential biomarker for biguanide sensitivity in cancer.

Enhanced Myc activity leads to overexpression of miR-17∼92, which in turn lowers LKB1 expression. This leads to enhanced tumor growth associated with increased glycolysis and oxidative phosphorylation, but it also reduces the capacity of lymphoma cells to activate the LKB1-AMPK system to overcome the energetic stress imposed by biguanides. Another important function of LKB1-AMPK signaling is the surveillance of mitochondrial integrity *(37)*. The potential impact of *miR-17∼92* on mitochondrial homeostasis and changes in response to biguanide treatment remains to be determined.

Our results address gaps in knowledge related to the mechanism by which LKB1 loss induces biguanide sensitivity. Previous work indicates that phenformin treatment increases ROS levels in cells with reduced LKB1 expression *(8)*, leading to oxidative damage and preferential cell death of LKB1-null tumor cells. We show here that glucose- and glutamine-dependent fueling of the TCA cycle was inhibited in *miR-17∼92*-expressing lymphoma cells responding to biguanide treatment. This fueling is a key node for the production of reducing equivalents for ATP production and biosynthetic intermediates for growth (e.g., citrate and aspartate). Our data are consistent with observations that reducing mitochondrial aspartate production in tumor cells can impair cell viability, due in part to diminished purine and pyrimidine biosynthesis *(38, 39)*. Reduced biosynthetic metabolism, combined with reduced mitochondrial NADH production and an inability to activate AMPK signaling, appear to sensitize *miR-17∼92*-expressing lymphoma to metabolic stress. Consistent with this, biguanide treatment—particularly IM156—slowed lymphoma cell growth in vitro and extended the lifespan of tumor-bearing mice.

There has been increased interest in the clinical use of biguanides for cancer therapy. Encouraging preclinical data *(17)* led to more than 100 clinical trials of metformin for indications in oncology *(21)*, but early results have been disappointing. Lack of clinical benefit of metformin in trials reported to date does not necessarily imply that biguanides as a class have no activity. Pharmacokinetic factors may account at least in part for the discrepancies between preclinical models and clinical trials with respect to antineoplastic activity of metformin. The hydrophilic nature of metformin dictates that cellular uptake is dependent on organic cation 1 (OCT1) transporters before gaining access to mitochondria *(29)*. This inherently limits the action of metformin to those cells expressing OCT1, which can vary between tumors *(40)*. The biguanide phenformin displays greater bioavailability and anti-neoplastic activity than metformin *(29, 40, 41)*, but its clinical use has been limited due to the risk of lactic acidosis, especially in patients with compromised renal or liver function *(42)*. These results justify investigation of novel biguanides with better pharmacokinetic/pharmacodynamic (PK/PD) properties than metformin in neoplastic tissue as a strategy for OXPHOS inhibition for cancer treatment. We identified IM156 from a library of >1,000 biguanides as a potential anti-cancer drug candidate *(43)*. Our data indicate that IM156 is a more potent OXPHOS inhibitor than metformin or phenformin. IM156 may have advantages over metformin as it is a more potent, hydrophilic biguanide that we hypothesize is more bioavailable in neoplastic tissues. IM156 is presently being evaluated in a phase I clinical trial (NCT 032 72256).

Our research adds to prior evidence *(8, 44)* that clinical trials of biguanides or other OXPHOS inhibitors are likely to show clinical activity if evaluated in patients with cancers that are particularly sensitive to energetic stress, such as those with elevated expression of *Myc* or *miR-17∼92* and/or decreased function of LKB1. Although LKB1 inactivation is relatively common in NSCLC *(7)*, it is not common amongst other malignancies. In contrast, *Myc* is among the most commonly dysregulated oncogenes in both hematologic malignancies and solid tumors. Because of the paucity of viable treatments targeting *Myc*-driven cancers, linking the downstream *Myc* target *miR-17∼92* to inhibition of LKB1 may identify a cohort of patients without LKB1 mutations who are particularly sensitive to biguanide treatment, opening new avenues for the treatment of these cancers.

## Materials and Methods

### Synthesis of IM156

IM156 was synthesized as previously reported *(45)*. In brief, pyrrolidine was dissolved in butanol before concentrated hydrochloric acid was added and stirred for 30 min at 0°C. Sodium dicyanamide was then added and stirred for 24 h under reflux. After completion of the reaction was confirmed, the adduct—N1-pyrollidine cyanoguanidine—was purified. Next, phenylamine was dissolved in butanol before concentrated hydrochloric acid was added and stirred for 30 min at room temperature. The N1-pyrollidine cyanoguanidine prepared above was then added and stirred for 6 h under reflux. This mixture was concentrated under reduced pressure then dissolved in a 6 N hydrochloric acid/methanol solution before adding ethyl acetate. The precipitate— IM156A HCl—was filtered and dried under reduced pressure. IM156.HCl can be converted to free form by adding sodium hydroxide. The free form can be converted to the acetate salt form— IM156A—by adding acetic acid.

### Cell lines, DNA constructs, and cell culture

The generation of Eµ-*Myc Cre-ERT2*^+^; *miR-17∼92*^*fl/fl*^ lymphoma cells has been described previously *(31)*. Deletion of *miR-17∼92* was achieved by culturing Eµ-*Myc Cre-ERT2*^+^;*miR-17∼92*^*fl/fl*^ cells with 250 nM 4-OHT for four days *(31)*, followed by subcloning 4-OHT-treated cells to isolate cells deficient for *miR-17∼92*. Eµ-*Myc* cells were cultured on a layer of irradiated *Ink4a*-null MEF feeder cells in DMEM and IMDM medium (50:50 mix) supplemented with 10% fetal bovine serum (FBS), 20000 U/mL penicillin, 7 mM streptomycin, 2 mM glutamine, and β-mercaptoethanol. Raji cells were cultured in RPMI medium supplemented with 10% FBS, 20000 U/mL penicillin, 7 mM streptomycin, and 2 mM glutamine. Cells were grown at 37°C in a humidified atmosphere supplemented with 5% (v/v) CO_2_.

Retroviral-mediated gene transfer into lymphoma cells was conducted as previously described *(46)*. Briefly, lymphoma cells were transduced via spin infection, followed by culture in 4 µg/mL puromycin for four days, and subsequent subcloning by limiting dilution. *miR-17∼92* constructs have been previously described *(31)*. Knockdown of *Stk11* via shRNA (sequence: 5’-AGGTCAAGATCCTCAAGAAGAA -3’) has been described *(28)*.

### Cell proliferation and viability assays

Cells were seeded at a density of 1 × 10^5^ cells/mL in 3.5 cm dishes, and cell counts determined via trypan blue exclusion using a TC20 Automated Cell Counter (Biorad). For viability measurements, cells were stained with Fixable Viability Dye eFluor 780 (eBioscience), and analyzed using a Gallios flow cytometer (Beckman Coulter, Fullerton, CA) and FlowJo software (Tree Star, Ashland, OR).

### Seahorse XF96 Respirometry and metabolic assays

Cellular oxygen consumption rate (OCR) and extracellular acidification rate (ECAR) were determined using an XF96 Extracellular Flux Analyzer (Seahorse Bioscience) using established protocols *(4, 47)*. In brief, 7.5 × 10^4^ lymphoma cells were plated per well of an XF96 Seahorse plate in 140 μL of unbuffered DMEM containing 25 mM glucose and 2 mM glutamine, followed by centrifugation at 500xg for five minutes. Seahorse plates were pre-coated with poly-D-lysine (Sigma-Aldrich) to enhance cell adherence. XF assays consisted of sequential mix (3 min), pause (3 min), and measurement (5 min) cycles, allowing for determination of OCR and ECAR every 8 min. Following four baseline measurements, 20 µL of untreated media, phenformin, or IM156 were injected into respective wells, and OCR and ECAR tracked over time.

### GC-MS analysis of ^13^C-labelled metabolites

Cellular metabolites were extracted and analyzed by GC-MS using previously described protocols *(4, 26, 48)*. Eµ-*Myc* cells (3-5 × 10^6^ per 3.5 cm dish) were incubated for 2 hours in untreated, 100 µM phenformin, or 10 µL IM156 medium containing 10% dialyzed FBS and [^13^C]-glutamine (Cambridge Isotope Laboratories). Cells were washed twice with normal saline, then lysed in ice-cold 80% methanol and sonicated. For GC-MS analysis, D-myristic acid (750 ng/sample) was added to metabolite extracts as an internal standard prior to drying samples by vacuum centrifugation with sample temperature controlled at -4 °C (LabConco). Dried extracts were dissolved in 30 µL methoxyamine hydrochloride (10 mg/ml) in pyridine and derivatized as tert-butyldimethylsilyl (TBDMS) esters using 70 µL N-(*tert*-butyldimethylsilyl)-N-methyltrifluoroacetamide (MTBSTFA). An Agilent 5975C GC-MS equipped with a DB-5MS+DG (30 m × 250 µm × 0.25 µm) capillary column (Agilent J&W, Santa Clara, CA, USA) was used for all GC-MS experiments, and data collected by electron impact set at 70 eV. A total of 1 μL of derivatized sample was injected per run in splitless mode with inlet temperature set to 280°C, using helium as a carrier gas with a flow rate of 1.5512 mL/min (rate at which D27-myristic acid elutes at 17.94 min). The quadrupole was set at 150°C and the GC/MS interface at 285°C. The oven program for all metabolite analyses started at 60°C held for 1 min, then increasing at a rate of 10°C/min until 320°C. Bake-out was at 320°C for 10 min. Sample data were acquired in scan mode (1–600 m/z).*(48)* Mass isotopomer distribution for TCA cycle intermediates was determined using a custom algorithm developed at McGill University *(48)*. After correction for natural ^13^C abundances, a comparison was made between non-labeled (^12^C) and ^13^C-labeled abundances for each metabolite. Metabolite abundance was expressed relative to the internal standard (D-myristic acid) and normalized to cell number.

### Immunoblotting and Quantitative Real-Time PCR

Lymphoma cell lines were subjected to SDS-PAGE and immunoblotting using CHAPS and AMPK lysis buffers as previously described *(46)*. Primary antibodies against β-actin, 4EBP (total, phospho-T36/47, and phospho-S65), rS6 (total and p S235/236), Raptor (total and pS792), and AMPKα (total and phospho-T172) were obtained from Cell Signaling Technology (Danvers, MA). Primary antibody against LKB1 (Ley 37D/G6) was obtained from Santa Cruz Biotechnology (Dallas, TX, USA). For qPCR quantification of mature miRNAs, Qiazol was used to isolate RNA, miRNEasy Mini kit was used to purify miRNAs and total mRNA, and cDNA wassynthesized using the miScript II RT kit (Qiagen). Quantitative PCR was performed using the SensiFAST SYBR Hi-ROX kit (Bioline) and an AriaMX Real Time PCR system (Agilent Technologies). miScript primer assays (Qiagen) were used to detect mature miRNAs of the *miR-17∼92* cluster, with miRNA expression normalized relative to U6 RNA levels.

### Tumor xenograft assays

Lymphoma cells were resuspended in HBSS at a concentration of 5 × 10^6^ cells/mL, and 10^6^ cells/200 μL were injected intravenously into CD-1 nude mice (Charles River). Water bottles carrying 1.2% sucralose, 0.9 mg/mL phenformin +1.2% sucralose, or 0.8 mg/mL IM156 + 1.2% sucralose were provided for ad lib consumption. Mice were tracked until clinical displays of disease, such as weight loss and poverty of movement, at which point mice were euthanized.

## Statistical Analysis

Statistics were determined using paired Student’s t-test, ANOVA, or Log-rank (Mantel– Cox) using Prism software (GraphPad) unless otherwise stated. Data are calculated as the mean ± SEM for biological triplicates, and the mean ± SD for technical replicates unless otherwise stated. Statistical significance is represented in figures by: *, *p<0.05*; **, *p<0.01*; ***, *p<0.001*

## Supplementary Materials Index

Fig. S1, related to Fig. 2. *miR-17∼92* sensitizes lymphoma cells to apoptosis by biguanides

Fig. S2, related to Fig. 6. *miR-17/20* expression correlates with biguanide sensitivity in human lymphoma cells

Table S1, related to Fig. 6. EC_50_ and *miR17/20* expression in human lymphoma cell lines

## Acknowledgments

We thank members of the Jones laboratory and colleagues at ImmunoMet Therapeutics for advice regarding the manuscript. We acknowledge technical assistance from the Metabolomics and Flow Cytometry Core facilities at the VAI and McGill/GCRC.

## Author contributions

Experimental design and execution was conducted by SI, AG, AOD, GB, TFD, MNP, and RGJ. Essential reagents were provided by RL, MDM, NAJ, and SY. Bioinformatics and data analysis was conducted by SI, RDS, and RGJ. Data interpretation was performed by SI, AG, AOD, GB, DA, RDS, TFD, and RGJ. The manuscript was written and edited by SI, KSW, MSR, SY, MNP, and RGJ.

## Data and materials availability

All data associated with this study are available in the main text or the supplementary materials.

## Supplementary Materials

**Fig. S1.**
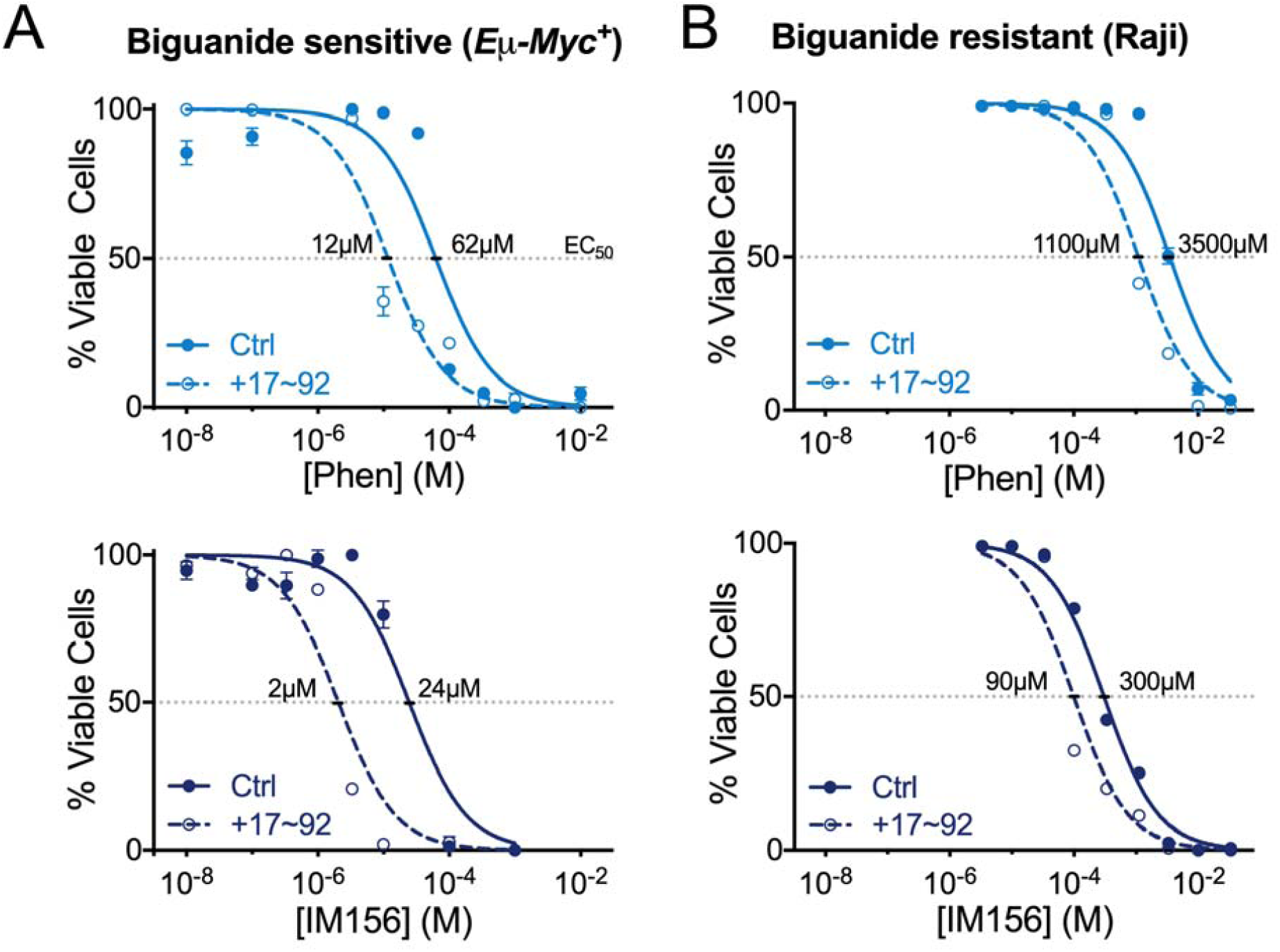
related to Fig. 2. *miR-17∼92* sensitizes lymphoma cells to apoptosis by biguanides. **(A–B)** Viability of biguanide sensitive Eµ-*Myc*^+^ lymphoma cells (A) or biguanide-resistant Raji cells (B) expressing control (Ctrl, closed circle/solid) or miR-17∼92 (+17∼92, open circle/dashed) vectors following 48 h treatment with indicated doses of phenformin (top) or IM156 (bottom). EC_50_ for each compound is indicated. Data represent mean ± SEM for biological replicates (n = 3 per drug per concentration). Statistics for all figures are as follows: *, *p<0.05*; **, *p<0.01*; ***, *p<0.001*.

**Fig. S2.**
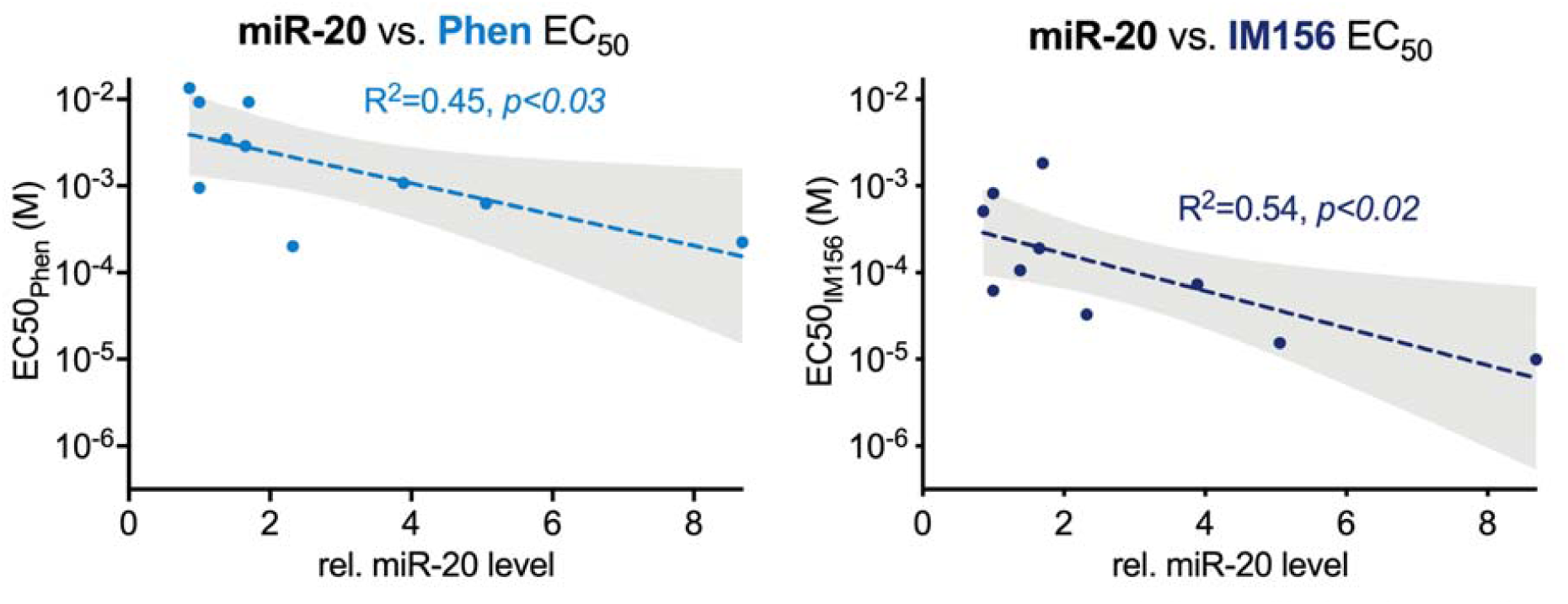
related to Fig. 6. *miR-17/20* expression correlates with biguanide sensitivity in human lymphoma cells. EC_50_ for phenformin (Phen, left) and IM156 (right) correlated to the relative levels of mature miR-20 transcript for ten human lymphoma cell lines. Linear regression (dotted line) is shown for both drugs with shaded 95% confidence interval. Statistics for all figures are as follows: *, *p<0.05*; **, *p<0.01*; ***, *p<0.001*.

**Table S1.** related to Fig. 6. EC_50_ and *miR17/20* expression in human lymphoma cell lines

|  | Relative expression |  |  | EC<sub>50</sub> |  |
| --- | --- | --- | --- | --- | --- |
|  | pri-miR-17 | miR-17 | miR-20 | Phen | IM156 |
| *SU-DHL-4* | 1 | 1 | 1 | 0.009283 | 0.0008229 |
| *OCI-Ly-7* | 8.49 | 1.47 | 2.32 | 0.0002009 | 0.00003293 |
| *Jeko-1* | 6.10 | 1.63 | 3.89 | 0.001075 | 0.00007331 |
| *Rec-1* | 11.08 | 2.04 | 5.06 | 0.0006291 | 0.00001547 |
| *Karpas 1718* | 12.69 | 4.21 | 8.69 | 0.0002254 | 0.000009992 |
| *OCI-Ly-1* | 1.07 | 1.05 | 1.38 | 0.003448 | 0.0001068 |
| *OCI-Ly-2* | 1.58 | 1.17 | 1.65 | 0.002892 | 0.00019 |
| *OCI-Ly-3* | 1.29 | 1.7 | 1 | 0.0009506 | 0.00006248 |
| *OCI-Ly-8* | 1.81 | 1.11 | 1.7 | 0.009283 | 0.001828 |
| *OCI-Ly-18* | 0.43 | 0.7 | 0.86 | 0.01349 | 0.0005098 |

